# Analysing the distribution of SARS-CoV-2 infections in schools: integrating model predictions with real world observations

**DOI:** 10.1101/2023.12.21.572736

**Authors:** Arnab Mukherjee, Sharmistha Mishra, Vijaya Kumar Murty, Swetaprovo Chaudhuri

## Abstract

School closures were used as strategies to mitigate transmission in the COVID-19 pandemic. Understanding the nature of SARS-CoV-2 outbreaks and the distribution of infections in classrooms could help inform targeted or ‘precision’ preventive measures and outbreak management in schools, in response to future pandemics. In this work, we derive an analytical model of Probability Density Function (PDF) of SARS-CoV-2 secondary infections and compare the model with infection data from all public schools in Ontario, Canada between September-December, 2021. The model accounts for major sources of variability in airborne transmission like viral load and dose-response (i.e., the human body’s response to pathogen exposure), air change rate, room dimension, and classroom occupancy. Comparisons between reported cases and the modeled PDF demonstrated the intrinsic overdispersed nature of the real-world and modeled distributions, but uncovered deviations stemming from an assumption of homogeneous spread within a classroom. The inclusion of near-field transmission effects resolved the discrepancy with improved quantitative agreement between the data and modeled distributions. This study provides a practical tool for predicting the size of outbreaks from one index infection, in closed spaces such as schools, and could be applied to inform more focused mitigation measures.

**Author summary:** At the start of the COVID-19 pandemic, there was huge uncertainty around the risks of SARS-CoV-2 spread in classrooms. In the absence of early predictions surrounding classroom risks, many jurisdictions across countries closed in-person education. There is great interest in adopting a more ‘precision’ approach to better inform future interventions in the context of airborne virus risks. For this purpose, we need tools that can predict the probability of the size of outbreaks within classrooms along with the impact of interventions including masks, better ventilation, and physical distancing by limiting the number of students per classroom. To this end, we have developed a robust but practical model that yields the probability of secondary infections stemming from index cases occurring within schools on a given day. During model development, the major underlying physical and biological factors that dictate the disease transmission process, both at long-range and close-range, have been accounted for. This enables our model to modify its predictions for different scenarios - and possibly allows its use beyond schools. Finally, the model’s predictive capability has been verified by comparing its outputs with publicly available data on SARS-CoV-2 diagnoses in Ontario public schools. To our knowledge, this is the first time an analytical model derived from mostly first principles describes real-world infection distributions, satisfactorily. The quantitative match between the theoretical prediction and real-world data offers the proposed model as a possible powerful tool for better-informed precision pandemic mitigation strategies in indoor environments like schools.

## Introduction

The public health response to SARS-CoV-2 in 2019 demonstrated a critical and growing need to anticipate the probability and size of outbreaks and the potential impact of mitigation measures. Models that can, with reasonable fidelity, predict the secondary infection distribution of cases in different scenarios, serve this critical need in pandemic and outbreak preparedness. Such predictive models would need to capture ‘superspreading’, a fundamental property of respiratory virus transmission – particularly in the context of aerosolized or airborne transmission. Superspreading events occur when the presence of one index case leads to infection of several individuals in a very short span of time. As a mean value description of outbreak size is incapable of capturing such events, Lloyd-Smith et al. [1] demonstrated the effectiveness of empirically fitted outbreak size distributions at modeling heterogeneity in disease spread dynamics. But, to retain predictive capabilities, models cannot rely only on infection data and instead would have to account for all the mechanistic parameters that uniquely define each case of an outbreak. These input parameters and their effects are rooted in physical, biological, and behavioral factors.

Recent studies detail the different aspects of SARS-CoV-2 transmission that need to be accounted for. The need for modeling airborne transmission is emphasized by Morawska et al. [2] and Allen et al. [3]. The review by Prather et al. [4] highlights the ability of aerosols to remain airborne for an extended period of time. Chaudhuri et al. [5, 6] showed that the viral load is a major contributor in determining the large variations in the number of secondary infections. These were complemented by Chen et al’s [7] observations of viral load heterogeneity being strongly tied to the overdispersed nature of SARS-CoV-2 infections, along with Bhavnani et al’s [8] findings of a direct correlation between higher index case viral loads and rising secondary infection counts. The alleviating effect of increased ventilation, incorporated through the air change rate parameter, on infection spread was thoroughly detailed by Thornton et al. [9], while Ricolfi et al. [10] studied this effect in schools and found that the inclusion of mechanical ventilation as opposed to natural ventilation drastically reduces the likelihood of infection. The importance of occupancy in infection spread was pointed out by Bazant and Bush [11]. Pioneering works by Haas [12–14] put forward the dose-response model which connects the virus properties to the human physiology and its response to the infectious dose. Schijven et al. [15] utilized the dose-response model to derive a risk assessment model for various expiratory events. Consolidating these findings, Chaudhuri et al. [5] proposed a model for the Probability Density Function (PDF) of secondary infections due to long-range airborne transmission, which relied on observations based on certain numerical results generated from cell-phone based occupancy data [16]. In this study, we systematically develop a model for the PDF of secondary infections from the equation governing the virion concentration at an indoor location, and couple its solution to the dose-response model governing pathogen-host interactions. The formulation is extended to an ensemble of locations to obtain a PDF for secondary infections due to long-range airborne transmission, ensuring an exclusively theoretical foundation. Additionally, this model is coupled with the contributions of short-range (or near-field) virus transmission to ensure that all the major sources of virus spread are captured. The final result is intended to be an analytical model capable of predicting outbreak sizes in real-world indoor locations while being able to adapt its solution to applied mitigation measures owing to its underlying theoretical foundation that captures the influence of such strategies.

One mitigation strategy applied during the pandemic has been widespread school closures, wherein teaching was moved to virtual classrooms. Emerging data suggest the move from in-person to virtual may have had a negative impact on students’ academic performance [17]. Perhaps, opportunities exist for more focused mitigation measures that include structural factors like enabling physical distancing with fewer students per classroom, ventilation, and more proximal measures like access and use of masks to offset the need to close schools in the event of a SARS-CoV-2 case in the classroom, or prior to detection of a case even when community transmission is high. In other words, with the ability to forecast the likelihood and magnitude of outbreaks in classrooms ahead of time, along with understanding the impact of various mitigation measures, making decisions about the advantages and disadvantages of school closures could be more well-informed. This opens an avenue for us to apply our model to judge its predictive capabilities in preparation for future outbreaks.

The primary objective of the present study is to derive an analytical model capable of predicting the real-world distribution of the number of secondary infections generated due to the spread of an airborne virus in schools and test its veracity through a comparison with real-world data. For this purpose, we use publicly available data on diagnosed SARS-CoV-2 cases from the public school system in Ontario, Canada, over different epochs in time. The modeling parameters chosen are based on existing literature, while also being guided by school data and mitigation strategies.

## Methods

The model is designed to predict the probability density function of the number of secondary airborne infections inside schools. The transmission process can be classified into two broad regimes: the virus traveling from the active index case to a susceptible individual; and the interaction between the virus and the human body. The former is mostly determined by physics-based factors while the latter depends on biological factors, primarily the dose-response of the human body.

First, we briefly review the dose-response model followed by the evolution equation of virus concentration in well-mixed situations i.e., virus concentration is homogeneous in space. Next, a discussion on the contribution of near-field virus concentration in assessing the overall outbreak sizes is provided. Coupling these with the distribution of the variables with the largest relative dispersions, the PDF of the number of secondary infections is derived with and without near-field effects.

For the convenience of the reader, a list of all symbols and their associated definitions are listed in Table 1 and Table 2 in S1 Text.

**Table 1.** Input Parameters for *α* and *α*_*j*_.

| Symbol | Definition | Value | Reference |
| --- | --- | --- | --- |
| $V$ | Classroom volume | 209 m<sup>3</sup> | Based on classroom size recommendation by a 2010 expert panel report [29] along with the assumption of classroom height $h = 3$ m |
| $\dot{V}_b$ | Inhalation rate | 105 cm<sup>3</sup>s<sup>-1</sup> | Inhalation rate is given as $\dot{V}_b = T_v W_a / \Delta$; $T_v = 7$ mL/kg is the average tidal volume per unit weight for children [30]; $W_a = 30$ kg is the assumed average weight of the susceptible individuals; $\Delta = 2$ s is the duration of inhalation based on the average breathing rate of 12 – 15 times per minute [31] |
| $Q_l$ | Ejected volume flow rate | $2.22 \times 10^{-6}$ mL/s | The exhalation rate of air for the index case is taken as 700 Lh<sup>-1</sup> or 194.4 cm<sup>3</sup>s<sup>-1</sup> [22]. This is combined with the aerosol size distribution for speaking [22] to obtain $Q_l$ as showcased by Chaudhuri et al. [5] |
| $F_{mask}$ | Mask filtration efficiency | 0.5 | Morais et al. [32] reported that commonly used homemade masks block 20% ($F_{mask} = 0.8$) to 60% ($F_{mask} = 0.4$) of all incoming aerosols, while N95 masks have much higher efficiency. We take an average value of 0.5 for our case |
| $\tau$ | Room-wide homogeneous virion field exposure duration | 6 hrs | Based on the total duration of classes in a school day [33] |
| $ACH$ | Air change rate | 2 hr<sup>-1</sup> | Reported by Swyers and King, CBC news article, 2021 [34] |
| $\beta_0$ | Wall-deposition parameter | 0.002 s<sup>-1</sup> | Refer to the calculations of Chaudhuri et al. [5] based on the study by Lai and Nazaroff [35] |
| $t_{1/2}$ | Virus half-life | 32.07 min | Based on the experiments of Dabisch et al. [36] at ASHRAE recommended indoor air conditions, using the DHS calculator [37] |
| $t_{tr}$ | Transition time to diffusion regime | 39 s | Defined as the time when cloud velocity is comparable to the ambient velocity fluctuations [38]. The final secondary infection PDF was found to be much less sensitive to $t_{tr}$ compared to most other parameters, and hence, it was approximated as the time when the cloud velocity becomes 1% of the cloud ejection velocity |
| $\tau_j$ | Near-field cloud exposure duration | 60 s | Assumed to be a short duration on average |

**Table 2.** Parameters defining the three cases computed in this study.

| Symbol | Definition | Default | Lower bound | Upper bound |
| --- | --- | --- | --- | --- |
| $\mathcal{T}$ | Total speaking duration of the index case in one school day | 2.4 hr | 1.2 hr | 3.6 hr |
| $\tau_j$ | Near-filed cloud exposure duration per person | 60 s | 30 s | 120 s |
| $t_{tr}$ | Puff-to-diffusion regime transition time (based on % of ejection velocity) | 39 s (1%) | 9 s (3%) | 100 s (0.05%) |
| $\mu_j$ | Mean of $\ln(N_j)$ – parameter governing the susceptible count that undergoes near-field transmission | 0.5 | 0.1 | 1.0 |
| $\sigma_j$ | Standard deviation of $\ln(N_j)$ – parameter governing the susceptible count that undergoes near-field transmission | 1 | 0.75 | 1.25 |

### Dose-response model

Quantitative risk assessment of infections is primarily performed based on two methods [18]. First among these is the well-known Wells-Riley [19, 20] approach that provided a relation connecting a quantum of infection to the probability of being infected. The simplicity of this relation allowed for application in various scenarios but it is unable to capture the underlying mechanisms governing the transmission. On the other hand, the second approach of dose-response models builds upon an analogous framework, while not being restricted by a hypothetical description for infectious dose i.e., quanta, therefore allowing these models to describe the infectious dose in terms of various biological and physical parameters that influence it. These models were first proposed by Haas [12] in the context of waterborne disease spread, but were also later applied in the case of SARS-CoV-2 [13, 21] as

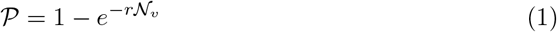

where the probability of infection is described by 𝒫, and 𝒩_*v*_ is the number of virions inhaled by a susceptible. The dose-response constant *r* was explained by Haas [13] as the inverse of the probability of a single virus surviving till it can trigger illness. The probability of infection *𝒫* has two interpretations - (1) if *𝒩*_*v*_ is the virion quantity inhaled by a person at a given location and time, then *𝒫* corresponds to that individual’s probability of getting infected from the inhaled virions; (2) if *𝒩*_*v*_ is the average infectious dose inhaled by a group of susceptibles, then 𝒫 represents the proportion of susceptibles that will become infected. The latter interpretation will be applied in the present model.

### Secondary infection count from the dose-response model

For the probability of infection 𝒫, we need to compute 𝒩_*v*_ in terms of known or computable quantities that can be obtained from existing school data and index case data. To that end, consider a classroom with one index case and multiple susceptible individuals. At any point in time, the susceptible individuals would be exposed to an airborne virion concentration field 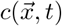. Here, a well-mixed room is assumed to invoke 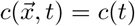 i.e., the virion concentration is homogeneous in space. If the index case possessing a viral load *ρ* continuously ejects mucosalivary fluid at a rate of *Q*(*t*) within a room of volume *V*, then the virion concentration *c* is governed by

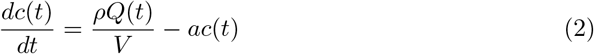

where the effects of air change rate *ACH*, wall deposition parameter *β*_0_, and virus half-life *t*_1*/*2_ appear through the loss parameter *a* = *ACH/*3600 + *β*_0_ + ln(2)*/t*_1*/*2_ [5]. The ventilation effect is assumed to be uniform here, though it is recognized that practically the aerosol flow can be anisotropic with a degree of directionality. The jet-puff model introduced later will partially address this. The volume flow rate *Q*(*t*) is assumed to have a constant value *Q*_*l*_ for the duration of the ejection event 𝒯, followed by a zero value for the remaining duration. The mucosalivary fluid ejection event under consideration is speaking, while breathing is neglected due to its comparatively (several orders of magnitude) lower contribution of ejected volume [22]. The solution of Eq. 2 takes the form

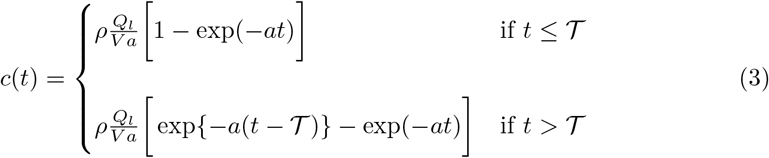

For this piecewise description of concentration *c*(*t*), the regime bounded by *t* ≤ 𝒯 marks the speaking duration of the index case. Beyond that, when *t >* 𝒯, the virion concentration decays. If the susceptible individuals are inhaling these virions at an average rate of 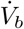 for a mean exposure duration of *τ*, the average number of virions inhaled by a single individual 𝒩_*v*_ can be written up to the leading order as

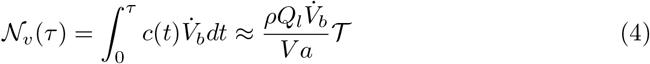

This leading order form was obtained through a series expansion of the result of the integral in Eq. 4 along with the application of a long exposure duration condition (*aτ* ≫ 1), which is easily satisfied in classroom scenarios. At this stage, Eq. 4 leaves us with a computable form for 𝒩_*v*_, which can now be substituted back in the dose-response model in Eq. 1 to obtain the probability of infection as

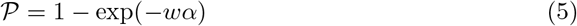

where

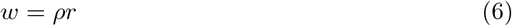

captures the primary sources of variability in 𝒫, while *α* is a constant that incorporates all other factors that influence the transmission process, and takes the form

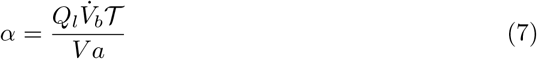

Observe that *α* admits mostly physics-based parameters within its description, whereas *w* is composed exclusively of biological parameters. This means that if the aerosol transmission physics changes it would be reflected through the constant *α*.

In Eq. 5, 𝒫 represents the proportion of secondary infections in a classroom with occupancy *n* and area *A*, from which the total number of secondary infections *Z* occurring there, under well-mixed conditions, is

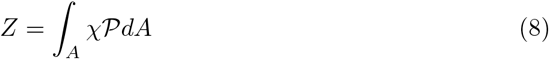

where *χ* = *n/A* is the population density in the classroom. The number of secondary infections *Z* can be expressed through a more generalized term which is independent of occupancy – the secondary attack rate 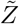, defined as the ratio of the number of secondary infections to the number of susceptibles at a location, such that 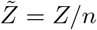. Noting that an individual’s probability of infection is independent of their spatial location due to the well-mixed room assumption, using Eq. 8, 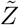 takes the form

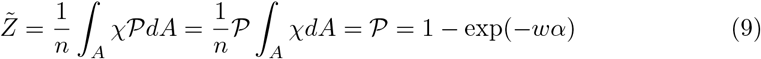

The invariance with occupancy makes the secondary attack rate a measure for the proportion of people infected, analogous to the probability 𝒫. The number of secondary infections due to one index case in a classroom can now be represented in terms of known or computable quantities as

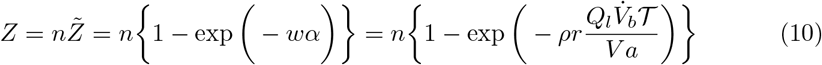

If the index case and the susceptible individuals are wearing masks with filtration efficiency *F*_*mask*_, it would modify the ejection and inhalation terms in Eq 10, now represented by *F*_*mask*_*Q*_*l*_ and 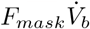 respectively.

Note that this entire derivation for long-ranged transmission has been performed for a classroom. But, the results we finally intend to obtain are for a school. How this long-range analysis for classrooms can be extended to schools will be explained in the Results section based on certain observations from real-world infection data.

### Modified secondary infection count through the inclusion of near-field transmission effects

Up to this point, the model developed only captures long-range airborne transmission within a classroom by invoking the well-mixed assumption. But in reality, there will be instances where the susceptible individuals will come into contact with the highly concentrated virion-laden cloud ejected by the index case before it diffuses. Even in scenarios where there are almost no infections through long-range transmission by virtue of the index case having a very low viral load, this cloud carrying a much higher virion concentration compared to the well-mixed case might still be capable of infecting those who come in contact with it. The concentration is higher since the ejected virions are localized in a smaller volume. To capture this effect, an equation for a modified secondary infection count *Z*_*m*_ is written

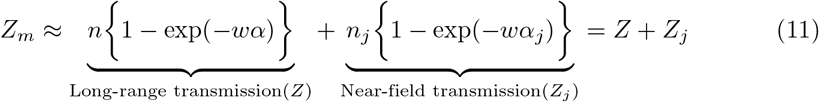

where *n* is the occupancy of the classroom that hosts the index case, and *n*_*j*_ is the total number of people in the school exposed to near-field transmission. Here, the total number of secondary infections has been modified simply through the addition of a near-field contribution *Z*_*j*_ which mathematically varies from the existing *Z* by having a modified occupancy *n*_*j*_ and a modified constant *α*_*j*_ that reflects the difference in the underlying physics of near-field transmission compared to the long-range route. Note that susceptibles counted in *n* that undergo long-range transmission may also get counted again in *n*_*j*_, which would appear to occasionally overestimate the total infection count. A detailed mathematical proof showcasing why such an overestimation will not occur is provided in the S2 Text.

The question remains about how to obtain the modified constant *α*_*j*_ that describes the near-field transmission physics. This is introduced through a simple jet-puff model for the ejected cloud that provides us with the cloud volume at any time *t*. An analogous assumption to that of the long-range model is applied i.e., the virion concentration within the cloud is spatially homogeneous. This enables us to use the same approach as the long-range model and solve a concentration equation for virions within the cloud up to its transition to a diffusion-dominated flow, assuming negligible concentration losses due to the transitory nature of the cloud, and subsequently obtain

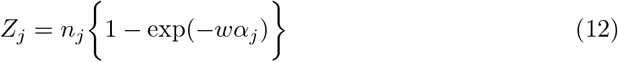

where the modified transmission physics is admitted through the constant *α*_*j*_ as

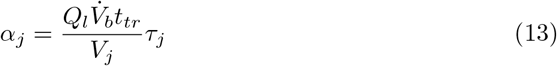

In Eq. 13, the transition time to diffusion dominated flow *t*_*tr*_, the volume of the cloud *V*_*j*_, and the average exposure duration to the cloud *τ*_*j*_ are new parameters unique to the near-field model. Here, *t*_*tr*_ and *τ*_*j*_ are input quantities whose values are discussed later, whereas the cloud volume *V*_*j*_ is calculated using a simple jet/puff model in S3 Text.

### Analytical PDF of total secondary infection count

At this stage, all the governing equations (Eq. 10, 11, and 12) required to describe the transmission process within a single location have been derived, and hence we have a closed system of equations. However, the goal is to model the variability of secondary infection count across different locations. This can be captured through a probability density function (PDF). The algorithm to obtain said PDF is shown in the flowchart provided in Fig. 1. Before proceeding, let us define the following pairs of quantities and their corresponding sample space variables: (*ρ, ρ*_*v*_), (*r, r*_*v*_), (*w, w*_*v*_), (*n, N*), (*n*_*j*_, *N*_*j*_) 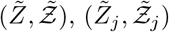, (*Z, Ƶ*), (*Z*_*j*_, *Ƶ*_*j*_), and (*Z*_*m*_, *Ƶ*_*m*_).

**Fig 1.**
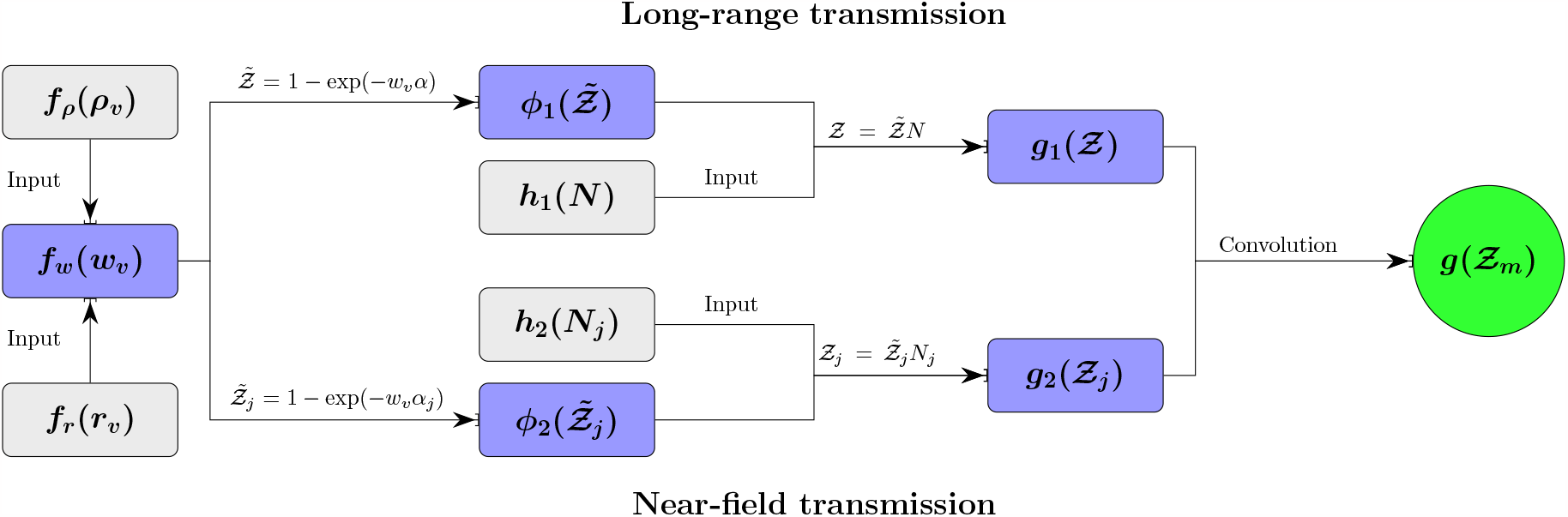
Algorithm for deriving the analytical PDF of total secondary infection count. The input distributions for viral load *ρ*_*v*_ and dose-response constant *r*_*v*_ are used to compute the PDF of *w*_*v*_ : *f*_*w*_(*w*_*v*_). This is applied in both halves of the transmission process – long-range and near-field, to find t_1_he PDF of the corresponding secondary attack rates, 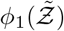 and 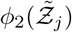. These coupled with their respective input occupancy distributions, *h*_1_(*N*) and *h*_2_(*N*_*j*_), gets us the PDF of secondary infections for long-range transmission *g*_1_(*Ƶ)* and near-field transmission *g*_2_(*Ƶ*_*j*_). A final convolution operation on these distributions outputs the total secondary infection count PDF, *g* (*Ƶ*_*m*_)

As shown in Fig. 1, we start from the input distributions for the most dispersive quantities – the viral load and dose-response constant, given by *f*_*ρ*_(*ρ*_*v*_) and *f*_*r*_(*r*_*v*_). These are used to compute the PDF of *w*_*v*_, given by *f*_*w*_(*w*_*v*_), which attains a lognormal form as derived later in the section detailing the inputs to our model.

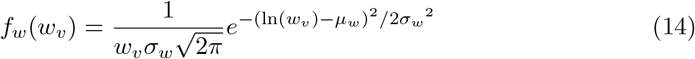

Here, *μ*_*w*_ and *σ*_*w*_ are the mean and standard deviation of ln(*w*_*v*_), respectively. This distribution is utilized in both long-range and near-field transmission routes without change because as stated before, the transmission physics is embedded in the constants *α* and *α*_*j*_. Mathematically, the PDF formulations for both routes are analogous, and hence, in the remainder of the section, we will only describe the long-range PDF in detail with the understanding that the same process can be followed for near-field transmission.

First, for long-range transmission, the secondary attack rate described by Eq. 9 is re-written as

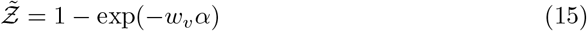

where *α* is a constant, and *w*_*v*_ has a distribution *f*_*w*_(*w*_*v*_). The justification behind assigning *α* an ensemble-averaged constant value instead of a distribution is discussed later. In Eq. 15, we observe a functional dependency of 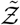 on *w*_*v*_. Therefore, the PDF of 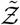, given by 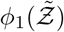, depends on *f*_*w*_(*w*_*v*_) as

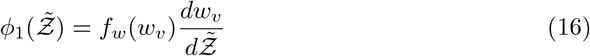

Rewriting 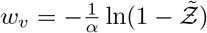 based on a re-arrangement of Eq. 15, we get

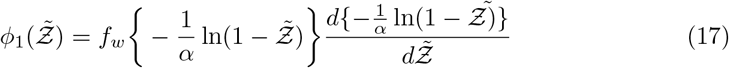

Substituting the lognormal form of *f*_*w*_(*w*_*v*_) in Eq. 17, 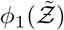 can be expressed as

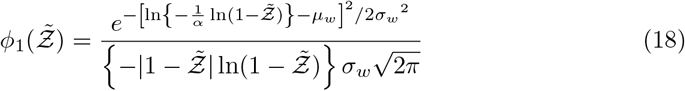

Now, to find the PDF of long-range secondary infection count *g*_1_(*Ƶ*), we need to combine 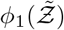 with the input PDF of classroom occupancy *h*_1_(*N*). This is achieved by employing the relation for finding the PDF of the product of two independent random variables 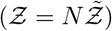 as

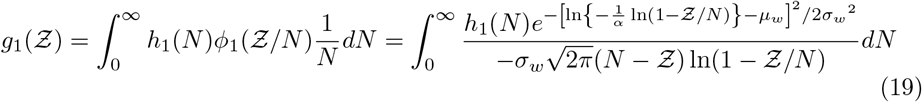

The exact procedure can now be repeated for near-field transmission to get an analogous PDF of near-field secondary infection count *g*_2_(*Ƶ*_*j*_).

Finally, a convolution operation is performed to obtain the PDF of the sum of two random variables, *Ƶ*_*m*_ = *Ƶ*+ *Ƶ*_*j*_; the PDF of net secondary infection count *Ƶ*_*m*_ is therefore computed as

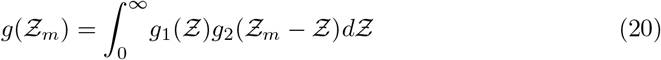

### Inputs to the analytical model

With the disease transmission model being completely defined through Eq 10, 12 and 20, we turn our focus towards the inputs that are required to solve the model and obtain the PDF of secondary infections. As observed before, the *Z* and *Z*_*j*_ equations (Eq. 10 and Eq. 12) require inputs to define three types of quantities: *w, α* (and *α*_*j*_), *n* (and *n*_*j*_). The primary sources of variability are embedded within *w* = *ρr* and hence, *ρ* and *r* will both admit individual distributions to account for this. On the contrary, *α* and *α*_*j*_ are composed of quantities that have comparatively negligible variation, and therefore, these quantities will all be assigned ensemble averaged values. Finally, *n* and *n*_*j*_ bring in the effect of occupancy through distributions of their own and is a highly relevant parameter since secondary infection count is linearly proportional to occupancy.

In the subsequent sections, we describe the input distributions for *ρ, r*, and *w*, followed by a section on all the ensemble-averaged parameters that generate *α* and *α*_*j*_, and end with a section describing occupancy distributions.

#### Probability density function for the viral load

Viral load *ρ* is a dominant factor that decides the shape of the *Ƶ* distribution, and hence its variability needs to be captured when providing it as an input parameter [5, 7]. The study by Yang et al. [23] collected viral load data for both asymptomatic and symptomatic individuals corresponding to the original variant of SARS-CoV-2, to generate a viral load PDF which showed a lognormal distribution of the form *f*_*ρ*_(*ρ*_*v*_; *μ*_*y*_ = 13.83, *σ*_*y*_ = 3.8) [5, 23], presented in Fig. 2(a) by a solid blue line.

**Fig 2.**
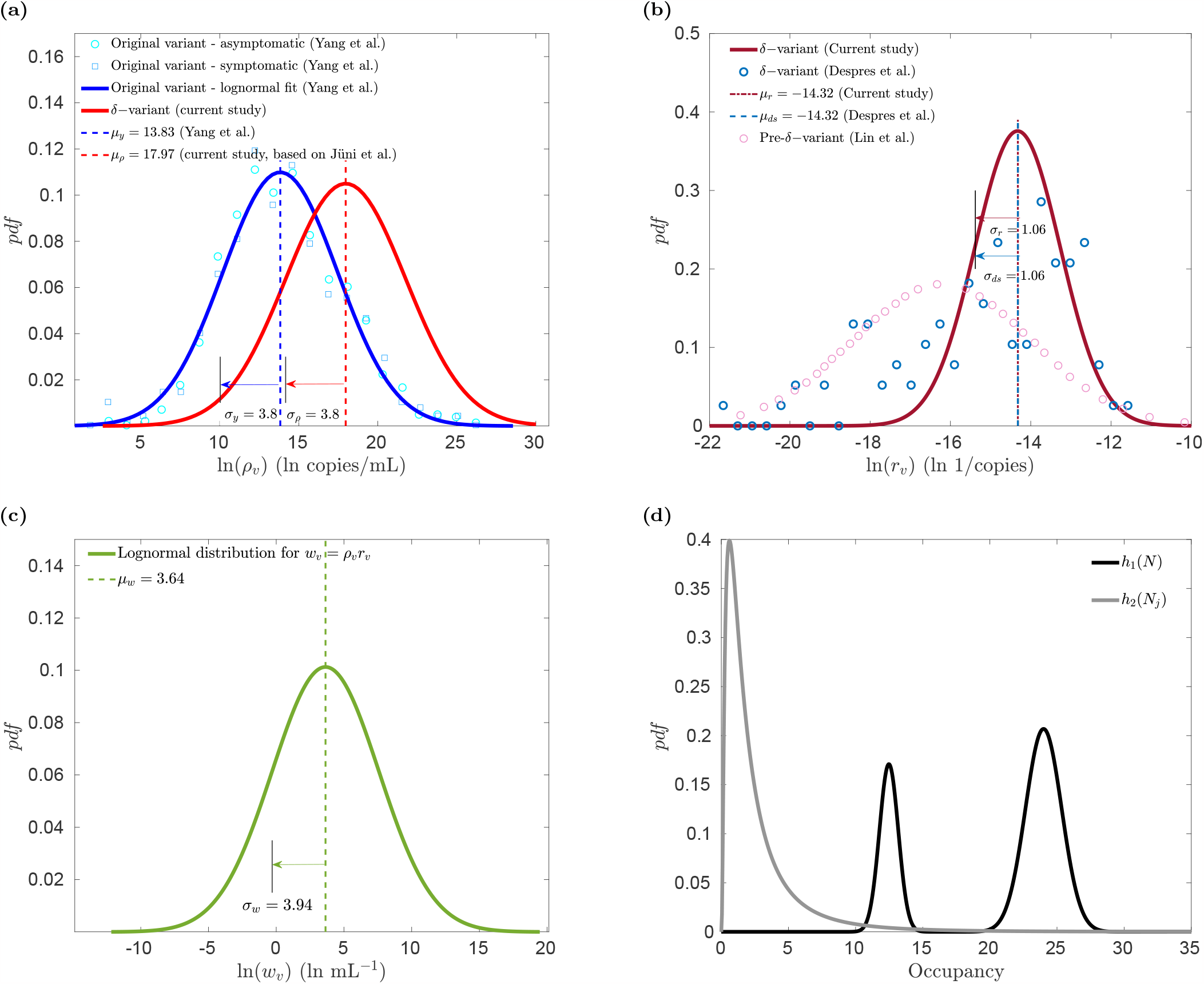
Probability density functions of input parameters. (a) Viral load *ρ*_*v*_ PDF for asymptomatic (circular markers) and symptomatic (square markers) population corresponding to the original SARS-CoV-2 variant [23] along with a lognormal fit (*μ*_*y*_ = 13.83, *σ*_*y*_ = 3.8) (solid blue line) for the data, compared to the *δ*-variant viral load lognormal distribution (*μ*_*ρ*_ = 17.97 [24], *σ*_*ρ*_ = 3.8 [23]) (solid red line), (b) Lognormal PDF of dose-response constant *r*_*v*_ for SARS-CoV-2 (pre-*δ*-variant) [28] (pink circles), compared to the PDF computed from *δ*-variant dataset [27] (blue circles), and a lognormal fit (dark red line) to this dataset ensuring equal mean and standard deviation (*μ*_*r*_ = *μ*_*ds*_ = −14.32, *σ*_*r*_ = *σ*_*ds*_ = 1.06), (c) Lognormal distribution for product of viral load and dose-response constant, *w*_*v*_ (*μ*_*w*_ = 3.64, *σ*_*w*_ = 3.94), (d) PDF of classroom occupancy *h*_1_(*N*) (black line) and PDF of the number of people exposed to near-field transmission, *h*_2_(*N*_*j*_) (grey line).

For the current study, we focus on a period between Sep-Dec 2021 when *δ*-variant was most dominant. Therefore, while retaining a similar lognormal form *f*_*ρ*_(*ρ*_*v*_; *μ*_*ρ*_, *σ*_*ρ*_), the distrtibution’s parameters need to be modified for the *δ*-variant. The *δ*-variant viral load vs. time plot from Jüni et al. [24] was used to calculate a time-averaged median viral load whose exponential gives *μ*_*ρ*_ = 17.97, and *σ*_*ρ*_ was assumed to be same as that obtained from Yang et al. [23]. The final *δ*-variant viral load distribution is also shown in Fig.2(a) by a solid red line.

#### Probability density function for the dose-response constant

The dose-response constant *r*, which has the units (RNA) ‘copies^−1^’, can be decomposed as

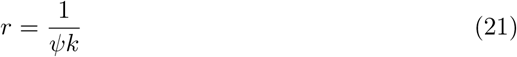

where *k* is an analogous dose-response constant parameter similar to *r*, and can be computed from literature, but generally presented in ‘Focus-forming units (FFU)’. To account for the difference in units between *r* and *k*, a quantity *ψ* is introduced whose inverse is often referred to as infectivity. In other words, *ψ* is a measure of the percentage of viral matter in droplets and aerosols that is capable of initiating infections. To find the PDF of *r* (corresponding sample space variable *r*_*v*_), the quantities *k* and *ψ* need to be determined.

For *k*, we refer to Killingley et al. [25], where they found that an infectious dose of 55 FFU of SARS-CoV-2 among volunteers showed 53% infection rate. Substituting these values in the dose-response model of Eq. 1 gives us *k* = 73 FFU. The literature regarding *k* values for SARS-CoV-2 are few and far between, and insufficient to obtain a distribution. However, assigning a constant *k* suffices because the primary variability in the dose-response constant appears from *ψ* which varies over several orders of magnitude [26–28].

For the PDF of *ψ*, we refer to a study by Lin et al. [28] that found *ψ* and therefore the dose-response constant to have a lognormal distribution, but not for the *δ*-variant. Therefore, using the dataset for *δ*-variant *ψ* from the study by Despres et al. [27], the dose-response constant’s mean and standard deviation were computed to assign the parameters of its lognormal distribution *f*_*r*_(*r*_*v*_; *μ*_*r*_, *σ*_*r*_) as

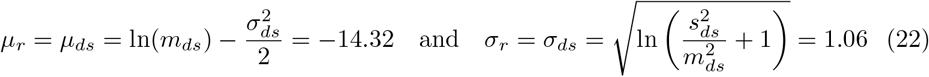

Here *m* is the mean of the data, and *s* is the corresponding standard deviation. The subscript ‘*ds*’ refers to the values being obtained using the data from Despres et al. [27]. The final PDF for *r*_*v*_ is displayed in Fig. 2(b) as a dark red line, compared to an approximate numerical distribution from Despres at al. [27] shown through blue circles. The pre-*δ*-variant distribution from Lin et al. [28] is also showcased in pink circles.

#### Probability density function for the product of viral load and dose-response constant

Since, *ρ*_*v*_ and *r*_*v*_ are both lognormally distributed, their product *w*_*v*_ = *ρ*_*v*_*r*_*v*_ also has a lognormal distribution with parameters *μ*_*w*_ = *μ*_*r*_ + *μ*_*ρ*_ = 3.64 and 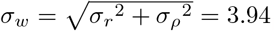. The PDF of *w*_*v*_ is shown in Fig. 2(c).

#### Description of the ensemble-averaged input parameters

All the ensemble-averaged parameters are embedded within *α* and *α*_*j*_, given by Eq. 7 and Eq. 13, and need to be properly defined as they introduce flow physics into the transmission model. Table 1 describes every parameter that is used to generate *α* and *α*_*j*_, along with their corresponding values, and how the values were computed. The question remains as to how we justify the parameters with *α* and *α*_*j*_ having ensemble-averaged values, as opposed to distributions like *ρ* and *r*. This can be answered if the dispersion for all these parameters across locations is calculated and compared. The quartile coefficient of dispersion for the *i*th parameter

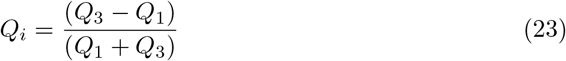

is employed for this purpose. Here, *Q*_1_ is the first quartile and *Q*_3_ is the third quartile of a given data set i.e., *Q*_*i*_ essentially provides a normalized measure for the spread of the data, while not being too sensitive to outliers in the dataset.

The quartile coefficient of dispersion *Q*_*i*_ for the different parameters in *w, α*, and *α*_*j*_ were calculated. For the quantities in *w* = *ρr, Q*_*ρ*_ = 0.98 and *Q*_*r*_ = 0.92 i.e., *Q*_*i*_ ≈ 1 for both, suggesting very high dispersion. In comparison, we have 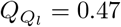 [5, 22, 39–42] and *Q*_*ACH*_ = 0.47 [5, 43], both much smaller than 1. Available values for the remaining quantities such as wall-deposition parameter [35], volume inhalation rates [22], virus half-life, classroom volumes, and others suggest significantly less variation. Therefore, *ρ* and *r* having much higher *Q*_*i*_ than every other quantity justifies assigning ensemble averaged values to every parameter in *α* and *α*_*j*_. Note that some parameters like ejection event duration, exposure duration to the ejected cloud, etc., do not have sufficient data in the literature to calculate dispersion. Hence, in the Results section, we have provided a range within which the PDF of secondary infections could lie to account for a certain degree of variation in such quantities.

#### Probabiliy density function of occupancy

The final step is to find an occupancy PDF that describes classroom populations. The present study has been carried out for the entire Ontario public education system, which as of 2020 − 21 data has a total of 2, 025, 265 students enrolled in 4, 833 schools, split into 1, 394, 040 elemendary students and 631, 225 secondary students [44]. This results in the average number of students per school being 419. Based on available data [45], the mean of the maximum classroom size is *μ*_1_ = 24 students with a standard deviation of *σ*_1_ = 1.35. A Gaussian distribution was fit based on these parameters as a first estimate for *h*_1_(*N*). But, this description neglects the effect of vaccination on the susceptible population distribution. We assume that elementary school students have not been vaccinated during the period of study. In contrast, secondary school students have a vaccination coverage of *η*_*cov*_ = 0.8 and a vaccination efficacy of *η*_*vac*_ = 0.6 [46]. The susceptible occupancy *n* is simply modified to *n*(1 − *η*_*vac*_*η*_*cov*_), and the modified Gaussian distribution has a mean of *μ*_2_ = 12.48 and a standard deviation of *σ*_2_ = 0.7. To incorporate both cases in a single model, a bimodal distribution *h*_1_(*N*) is chosen for occupancy, with a weighting factor of *p* = 0.3 (for secondary school students) based on the percentage of students in elementary and secondary schools. This results in a *h*_1_(*N*) distribution of the form

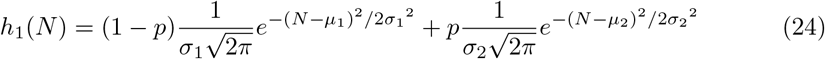

shown in Fig. 2(d) with a black line.

For the distribution of susceptibles exposed to near-field transmission *h*_2_(*N*_*j*_), we assume a long-tailed distribution to account for the likely large variation in the number of people interacting with the index case at close range. The simplest long-tailed distribution is chosen i.e., the lognormal. Its parameters were taken as *μ*_*j*_ = 0.5 and *σ*_*j*_ = 1 to ensure that the average number of near-field transmissions is low, which is expected because close-range interactions would be limited during a pandemic. Fig. 2(d) also showcases *h*_2_(*N*_*j*_) through a grey line.

## Results

The results presented correspond to two sources: real-life school infection data and modeled equations. Each source was used to obtain a probability density function for the total number of secondary infections occurring at a school within one school day.

### Analysis of reported school infection data

To generate the secondary infection PDF from actual cases, we draw on all reported SARS-CoV-2 diagnoses in 4,833 schools within the Ontario public school system. Ten distinct dates between 14 September 2021 and 13 December 2021 are chosen. The Ontario government official website [47] hosts the infection data reported by each school under its jurisdiction almost daily, which involves both student cases, and the grossly outnumbered staff cases that we therefore neglected. This data is sufficient to plot the PDF for secondary infections in schools within the province of Ontario for any date.

However, the generated PDF will be based on certain data points that correspond to scenarios where a school had no index cases and thus reported no new cases. Recall that our modeled PDF is only for scenarios where an index case is present. To exclude scenarios with no active index cases in a school, the following operations are performed.

Based on existing literature [24], we assume that an average of four days separate a susceptible getting infected in a school, and the school testing and reporting said infection. Fig. 3 shows the cumulative number of reported cases in Ontario in the two weeks before the day of infection [48], compared to those reported in Ontario public schools on the same dates. An immediate observation is that the number of school cases and Ontario cases are directly proportional, shown through the school infection to Ontario infection ratio in Fig. 3 being nearly constant over the three months under consideration. This allows us to assume that on the day of infection, the active index case proportion in the school population is equal to that in the Ontario population, given by *ϕ*_*i*_. Therefore, the average number of index cases per school *I*_*i*_ equals

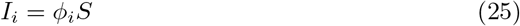

where *S* is the average student count in schools. Based on the data processed in this study, we note that all values of *I*_*i*_ *≤* 1. Therefore, assuming that index cases are spread evenly across *N*_*s*_ schools in Ontario, the total number of schools with at least one index case *N*_*i*_ on the day of infection is

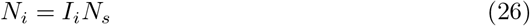

Next, we focus on the reported school infection data and note that they would comprise infections acquired in schools and those from outside schools. Our model does not account for the later contribution and hence it must be subtracted from the data. Thus, if *ϕ*_*j*_ is the active index case proportion in Ontario (and its schools) on the day the schools test and report for SARS-CoV-2, then the total number of outside contributions per school *I*_*j*_ is

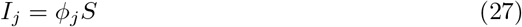

Therefore, if *Z*_data_ is the total infection count reported by a school, then the corrected infection count 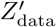 accounting for only in-school infections becomes

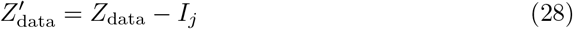

A histogram is then generated through proper binning of the 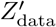 data set and then converted to a PDF using the relation

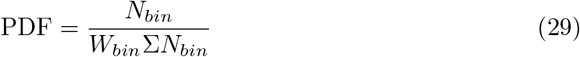

where *N*_*bin*_ corresponds to the number of elements in a bin i.e., the number of schools corresponding to a particular infection count, and *W*_*bin*_ = 1 is the bin width of the histogram. Note that the total bin element count Σ*N*_*bin*_ is not equal to the total number of schools *N*_*s*_, but instead equal to the total number of schools with at least one active index case on the day of infection, *N*_*i*_.

**Fig 3.**
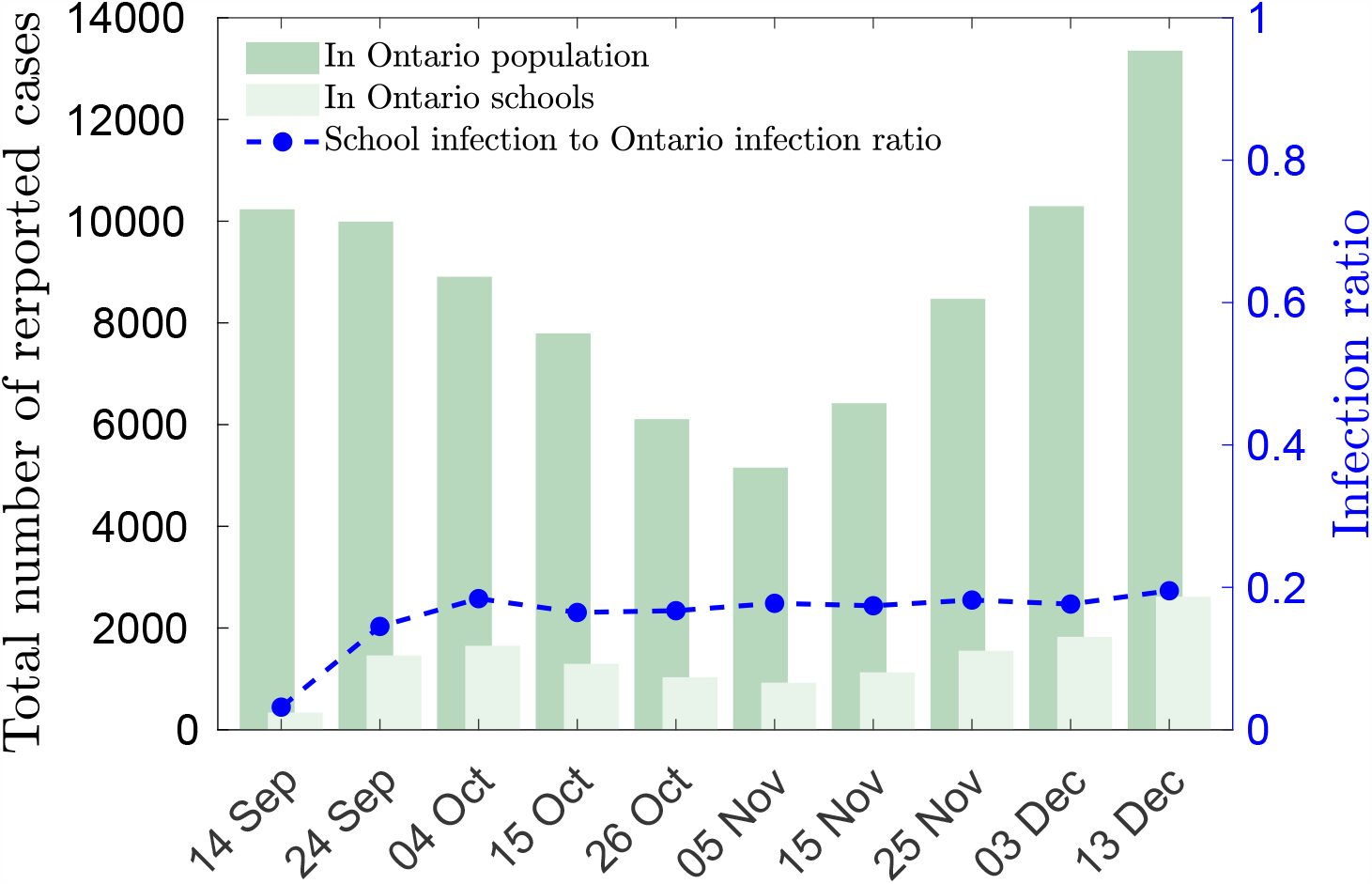
Reported SARS-CoV-2 cases in Ontario province and Ontario public schools. Bar charts for cumulative cases reported in Ontario in the two weeks before infection and cases reported in Ontario public schools for different dates are displayed. The blue curve corresponds to the ratio between cases reported in Ontario public schools to those reported in the entire province. Note that the total number of reported cases in Ontario corresponds to two weeks before the date of infection (assumed to be 4 days before infection was caught through testing) and not the date of reporting that is shown in the plot. All dates are for the year 2021.

We further clarify the data analysis with the following example. Consider the data reported in Ontario public schools on November 25, 2021. Based on usual SARS-CoV-2 incubation periods, the expected date of infection is taken as November 21, 2021. There were 8, 460 cases reported in the two weeks prior to this date. With Ontario’s population of 14 million, the index case percentage is *ϕ*_*i*_ = 0.06%. Superimposing this percentage on the *N*_*s*_ = 4, 833 Ontario public schools with *S* = 419 average students, we get *I*_*i*_ = 0.2532 index cases per school. This number would suggest that, on average, there are schools with no index cases i.e., 1 index case in approximately 4 schools. The modified school count would then become *N*_*i*_ = 0.2532 *×* 4, 833 = 1, 224 schools. The dates used for data analysis in this study are at least 8 − 10 days apart to ensure that the infection events are independent and there is no repeat counting of the same infections.

### Application of the analytical model in schools

As noted previously, our analytical model is derived for a single classroom scenario. But, in this study, we instead apply it to model SARS-CoV-2 spread in an entire school. We can justify this approach through the observation that, on average, a single classroom with one index case coupled with their near-field transmission contributions is sufficient to represent SARS-CoV-2 spread within a school. The chain of logic used to make this justification is shown in Fig. 4 and is further discussed below.

**Fig 4.**
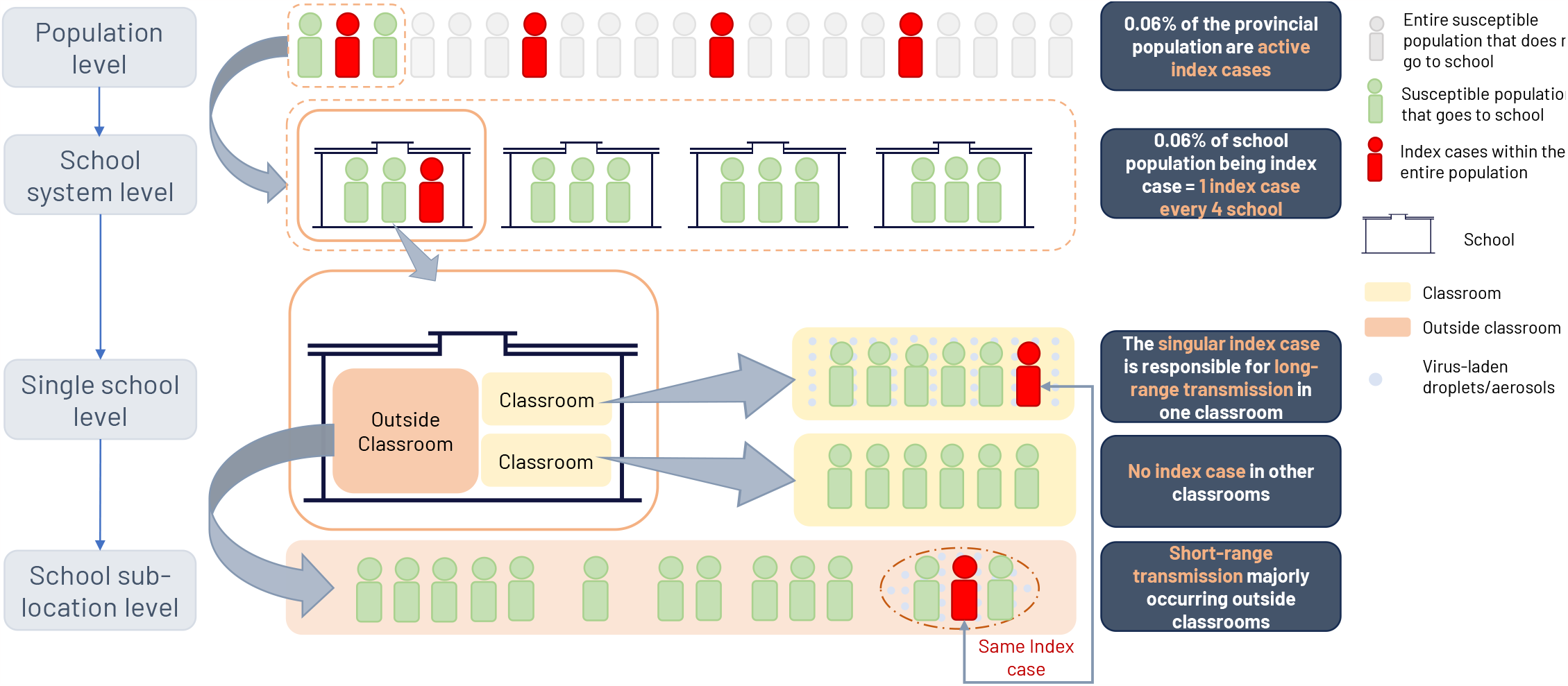
Representation of SARS-CoV-2 spread in a school through a single classroom coupled with outside-classroom effects. The schematic was built based on the example of SARS-CoV-2 infections in school on November 25, 2021. At the population level, individuals are classified into – active index cases who may or may not go to school, susceptibles that go to school, and susceptibles that do not go to school. Moving one level below, we have the entire school system consisting of the susceptible and index case population that goes to school. Here, data tells us that on average only one index case is active across multiple schools. Moving down another level, our focus is shifted to the school hosting that one active index case. Here, the index case divides their time inside a classroom or outside it. At the lowest level, we look at these individual sub-locations (classrooms, outside classrooms) where in one classroom the index case is the cause of long-range and near-field SARS-CoV-2 transmission, whereas every other classroom is devoid of active index cases and thus does not contribute to the disease spread within that school. Additionally, the index case spends time outside classrooms for a shorter duration in closer proximity to some individuals and thus spreads the virus through near-field transmission. Hence, the one classroom with the index case coupled with the index case’s outside classroom interactions can approximately represent SARS-CoV-2 spread within an entire school.

Referring back to the example of November 25, 2021, in the previous section, we focus on the population in Ontario – index cases and susceptibles, that go to school. For this date, we observe that on average 1 in every 4 school has an active index case i.e., 3 of 4 such schools do not host an index case and are neglected from our analysis. Next, we turn our attention to every such school that hosts an active index case. Here, the index case spends their time inside a classroom or outside it. As a large portion of their school day is spent within a classroom, this is where they become primarily a source for long-range airborne transmission. Their outside-classroom time, however, would involve small-duration interactions with others that are too short for long-range transmission, but instead contribute to the SARS-CoV-2 spread within that school through near-field transmissions. All other classrooms, which do not host active index cases, are non-contributors to the disease spread dynamics within that school.

These sets of observations showcase how a classroom with an active index case, coupled with that same index case’s near-field virus transmission contributions is sufficient to represent the average virus spread scenario in an entire school. Hence, our analytical model, which is able to account for these processes, can now be applied to schools.

### Comparison between modeled results and reported school data

The PDFs for the number of secondary infections *Z* (sample space variable *Ƶ*) at a school within a single school day, generated from the reported data for the 10 dates chosen, are shown in Fig. 5 using red symbols. All PDFs peak near zero, followed by a strong gradient that smoothens out into a long tail that highlights the overdispersion in SARS-CoV-2 transmission. This common behavior suggests that even though low infection counts are the norm in the presence of an active index case within a school, the existence of the occasional superspreading events described by the data points at large *Ƶ*, is a constant threat and a reminder for proactive mitigation measures. Most of these PDFs attain small peaks near *Ƶ* ≈ 10 and *Ƶ* ≈ 20, possibly pointing towards susceptible student populations being localized around that mark. The clustering of all the PDFs reveals a shared underlying trend likely described by the properties of a common virus strain (*δ*-variant) responsible for the infections during Sep-Dec 2021.

**Fig 5.**
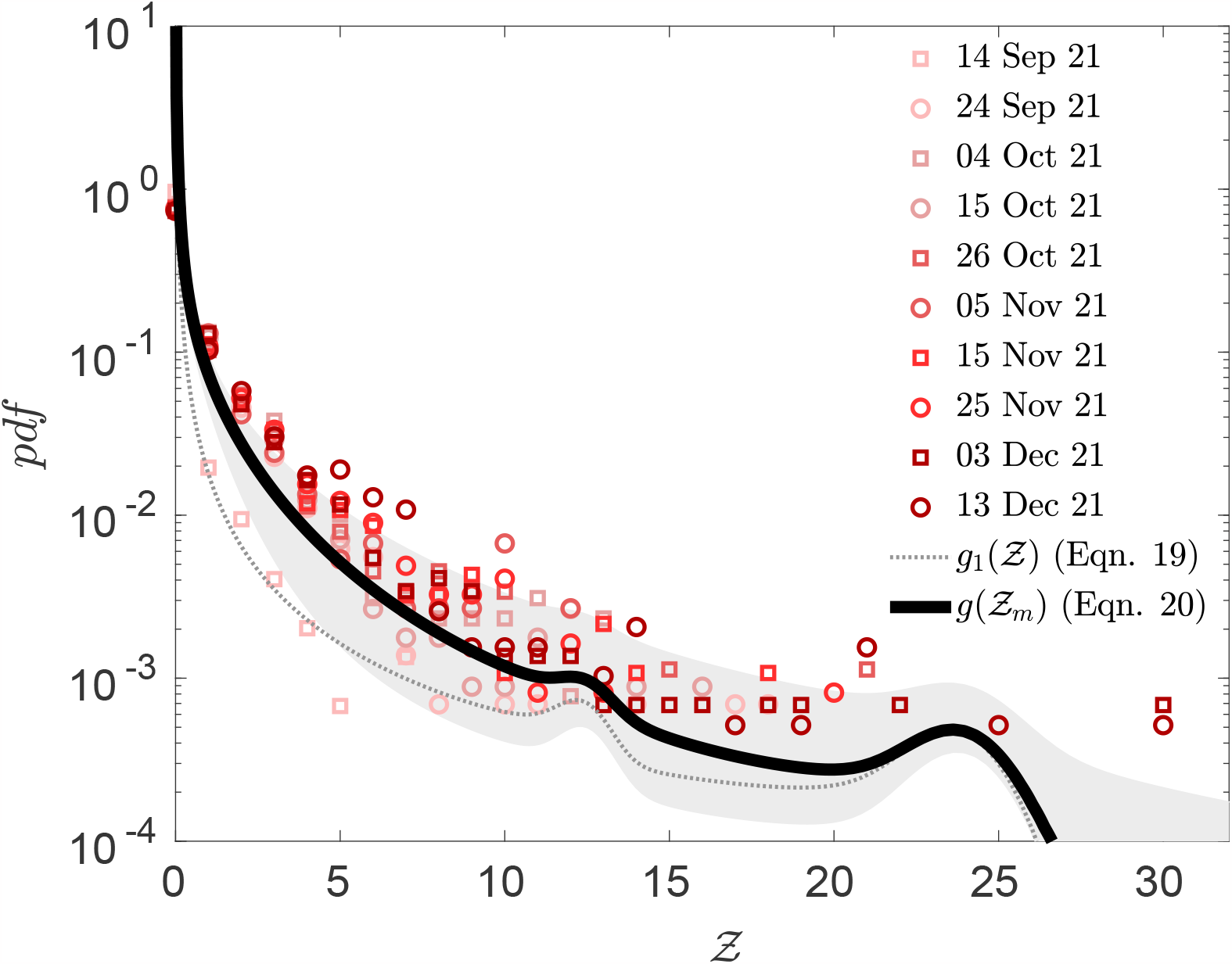
Model predictions for secondary infection distribution in schools compared with reported school infection data. Analytical PDF of secondary infections through long-range transmission *Ƶ* (Eq. 19) (thin dotted line) and with added near field transmission contribution *Ƶ*_*m*_ (Eq. 20) (solid black line), compared to reported infection data in Ontario public schools (red symbols). The shaded region provides an approximate measure for variation in *g*(*Ƶ*_*m*_) with a choice of different input values for certain parameters. The upper and lower bounds of the shaded region correspond to the cases listed in Table 2

In the remainder of this section, we will aim to test whether our current model can replicate such real-world data. To that end, we first employed the initial iteration of our model that solves only for the analytical PDF of total number of secondary infections, *g*_1_(*Ƶ*) (Eq. 19), due to long-range transmission from an active index case. The input parameters used to generate *g*_1_(*Ƶ*) account for both school-specific data and *δ*-variant specific biological quantities. The computed *g*_1_(*Ƶ*) PDF in Fig. 5 shown as a thin dotted line is compared to the reported data, clearly showcasing qualitative similarities while maintaining a degree of quantitative mismatch. In particular, the model notably underpredicts the PDF values at small *Ƶ ≤* 10, described by the much higher probability at zero, followed by a sharper gradient compared to the reported data. We hypothesized that this could be a limitation that arises from the assumption of homogeneous aerosol concentration, which cannot account for the increased number of infections in close proximity to the index case. The analytical PDF does successfully capture the long-tailed nature of the reported data, while also peaking twice in tandem with it, by virtue of the bimodal Gaussian nature of the classroom occupancy distribution.

However, its tail diverges from the reported data and falls off at high *Ƶ* (*Ƶ* ≥ 25). This is because the data sample size is limited, causing any unlikely high-valued samples to skew the corresponding probability, somewhat artificially. Additionally, the reported data PDFs have points that go beyond the maximum class size considered in the model, reflecting infections incurred outside classrooms, and is a limitation of representing the disease spread dynamics within a school exclusively through a classroom.

To ensure improved quantitative predictions from our model, a near-field effect was introduced – modeling the interactions of susceptibles with the localized, high virion density cloud ejected by the index case. This included coupling the well-mixed virion field with the ejected cloud represented by a simplified jet-puff model [6] to derive the PDF of secondary infections due to both long-range and near-field transmission *g*(*Ƶ*_*m*_) (Eq. 20). The modified PDF *g*(*Ƶ*_*m*_) is shown as a solid black line in Fig. 5 where an excellent quantitative match is immediately observable as *g*(*Ƶ*_*m*_) now passes through the cluster of red curves, and retaining the overdispersed nature of the older model. The magnitude of the gradient at low *Ƶ* regime is smaller, while the probability of zero infections has shifted to lower values, mimicking the reported data. These observations support our near-field contribution hypothesis and suggest that several of the previously zero infection scenarios now involve a non-zero near-field infection count. The near-field model also includes outside-classroom infections and removes the previous limitation of a classroom-specific description. The success of this updated model strongly emphasizes the need for aerosolized transmission models to capture both near-field and far-field mechanisms, while putting forward a practical tool that for the first time, to the best of our knowledge, reproduces real-world infection spread dynamics in schools from an exclusively theoretical foundation.

Finally, we revisit the fact that certain parameters used to compute the PDF of *Ƶ*_*m*_ have no real-life data source and have been ascribed an intuitive approximate value i.e., their assigned values likely have an error margin. To account for this possible variation, two more cases have been run by modifying these particular parameters – (1) for an approximate lower bound scenario corresponding to low virion concentration exposure compared to the default case, and (2) for an approximate upper bound scenario corresponding to high virion concentration exposure. Table 2 lists such input parameters for all three cases that have been used in this study. Note that these bounding parameters do not have extreme values (e.g. zero speaking duration for an index case in one school day), as these cases are not meant to account for outlier situations; those are already handled by the PDF and its long tail, but rather intended to highlight a deviation in the approximate average values assigned. The *g*(*Ƶ*_*m*_) curves for these cases enclose the shaded region in Fig. 5, providing an approximate range within which the secondary infection PDF could vary. The shaded region itself lies relatively close to the band of real-life data curves. Even among the quantities varied to obtain the bounds, not all influence *g*(*Ƶ*_*m*_) equally; quantities like transition time to diffusion barely affect the PDF, whereas others like *μ*_*j*_ and *σ*_*j*_ that govern the distribution of susceptibles exposed to near-field transmission have a much stronger influence. This gives us a glimpse into how mitigation measures, that influence other input parameters, and their effectiveness at curbing disease transmission will present themselves through the *g*(*Ƶ*_*m*_) plot. Variation in the input parameters due to possible mitigation measures such as better masks, higher ventilation, lowered occupancy, vaccination, etc., would either shift *g*(*Ƶ*_*m*_) to lower values or modify the shape of the PDF, and thus provide insight and justification behind opting for certain mitigation strategies over others, such that they allow for efficient school operations during future pandemics.

## Discussion

The present study puts forward a robust yet practical tool capable of predicting outbreak sizes of airborne diseases such as SARS-CoV-2 in schools or similar indoor locations, allowing for rapid informed implementation of ‘precision’ mitigation measures. This tool, in the form of an analytical model, was developed mostly from first principles and then examined against publicly available school infection data which guided further model improvements to boost its predictive capabilities.

Model development broadly involved coupling the dose-response function governing the human body’s response to incoming pathogens, with the physics of airborne transmission of said pathogens, to obtain the secondary infection count of an index case. First, we modeled the long-range transmission dynamics of the virus through a concentration equation in a well-mixed room, followed by expanding the model to an ensemble of rooms/locations, such that an analytical probability density function (PDF) of secondary infections could be derived. This stage saw the incorporation of several input parameters, among which the most dispersive ones – viral load and dose-response constant, were assigned distributions based on existing data. The choice of assigning an appropriate distribution instead of a mean value provided our model the flexibility of reacting to the biological processes that bring about such high dispersion in these quantities in the first place. The modeled PDF was then employed to compute the secondary infection distribution in Ontario public schools. The analytical nature of the predictive tool made this computation reasonably fast.

Our first set of comparisons between the modeled PDF based on homogeneous aerosol concentration and reported school data [47] displayed strong qualitative similarities plagued by a degree of quantitative mismatch. The similarities served as evidence of proper incorporation of the major governing factors of transmission, whereas the differences provided insights that led to the introduction of near-field transmission dynamics, modeled through a simple jet/puff model for the index case’s ejected virion cloud. A convolution operation on the individual outputs of the long-range and near-field models generated the net secondary infection PDF. Another set of comparisons with reported school data showed an excellent quantitative match, emphasizing the importance of the near-field model. At this stage, we have a model that not only is practical to use due to its analytical tractability but can predict real-world SARS-CoV-2 spread in schools. Apart from predicting future outbreak sizes for SARS-CoV-2 variants in schools, this has potential application toward other airborne disease outbreaks within a variety of indoor locations due to the mostly generalized nature of its underlying description.

Furthermore, the model can also revise its prediction based on the mitigation measure implemented. The final PDF generated, is sensitive to all its component parameters to varying degrees, each of which brings in some form of mitigation strategy. Reduction in inhaled or exhaled volume is affected by masks. Ventilation and ambient effects influence the virion concentration loss term in the formulation. Occupancy and room volume capture the role of population density in enclosed spaces. The physiological properties, viral load, and dose-response, would be a possible avenue to introduce the effects of developed immunity due to vaccinations and repeat infections. A change in any of these parameters in the model would pre-emptively inform the user about the efficacy of a proposed strategy and allow for the implementation of focused mitigation measures for future pandemics. This functionality allows our model to be a proactive predictive tool applicable to future airborne disease spread in schools and similar venues.

## Acknowledgments

AM acknowledges support from the Fields Institute for Research in Mathematical Sciences through their project Mathematics for Public Health and Variants of Concern sponsored by the Canadian Institutes of Health Research (Grant No. VS2-175577). SM acknowledges the Tier 2 Canada Research Chair in Mathematical Modeling and Program Science (Grant No. CRC-950–232643).

